# VP-Detector: A 3D convolutional neural network for automated macromolecule localization and classification in cryo-electron tomograms

**DOI:** 10.1101/2021.05.25.443703

**Authors:** Yu Hao, Biao Zhang, Xiaohua Wan, Rui Yan, Zhiyong Liu, Jintao Li, Shihua Zhang, Xuefeng Cui, Fa Zhang

## Abstract

**Motivation:** Cryo-electron tomography (Cryo-ET) with sub-tomogram averaging (STA) is indispensable when studying macromolecule structures and functions in their native environments. However, current tomographic reconstructions suffer the low signal-to-noise (SNR) ratio and the missing wedge artifacts. Hence, automatic and accurate macromolecule localization and classification become the bottleneck problem for structural determination by STA. Here, we propose a 3D multi-scale dense convolutional neural network (MSDNet) for voxel-wise annotations of tomograms. Weighted focal loss is adopted as a loss function to solve the class imbalance. The proposed network combines 3D hybrid dilated convolutions (HDC) and dense connectivity to ensure an accurate performance with relatively few trainable parameters. 3D HDC expands the receptive field without losing resolution or learning extra parameters. Dense connectivity facilitates the re-use of feature maps to generate fewer intermediate feature maps and trainable parameters. Then, we design a 3D MSDNet based approach for fully automatic macromolecule localization and classification, called VP-Detector (Voxel-wise Particle Detector). VP-Detector is efficient because classification performs on the pre-calculated coordinates instead of a sliding window.

**Results:** We evaluated the VP-Detector on simulated tomograms. Compared to the state-of-the-art methods, our method achieved a competitive performance on localization with the highest F1-score. We also demonstrated that the weighted focal loss improves the classification of hard classes. We trained the network on a part of training sets to prove the availability of training on relatively small datasets. Moreover, the experiment shows that VP-Detector has a fast particle detection speed, which costs less than 14 minutes on a test tomogram.

**Supplementary information:** Supplementary data are available at *Bioinformatics* online.

## 1 Introduction

Cryo-electron tomography (Cryo-ET) is currently the only imaging technique that allows 3D visualization of protein or protein complexes at molecular resolution in their native context. Subsequently, structures of interest presented as multiple noisy copies within a set of tomograms are computationally extracted, aligned, and averaged to yield higher resolution up to subnanometer resolution by a technique termed sub-tomogram averaging (STA) (Hutchings et al., 2018; Turonová et al., 2017; Schur et al., 2016). This technique provides biological insights into the interaction and function of structures imaged under close-to-life conditions. Localizing and classifying the macromolecules in cryo-electron tomogram is the first step for the STA process. The accurate particle localization and classification can benefit the subsequent alignment and averaging of subtomograms to improve the resolution of macromolecular structure.

However, particle localization and classification in cryo-electron tomogram is still challenging. One difficulty is the low signal-to-noise ratio (SNR) of tomographic reconstructions. Due to the beam-induced deformations in the biological sample, the electron dose applies to each tilt image very low, leading to a high amount of non-Gaussian noise for each image. The noise is further aggravated when the sample changes its thickness in the beam direction during tilting. Another trouble is the macromolecules of the same type are different from each other. It is because that each sub-tomogram loses the information in a wedge-shaped region in Fourier space and exhibits a deformation parallel to the direction of missing information in real space.

For the past decade, several software packages have been designed for Cryo-ET together with STA, such as PEET (Heumann et al., 2011), Eman2 (Chen et al. 2019), RELION (Bharat et al. 2016), Dynamo (Castaño-Díez et al. 2012; Castaño-Díez et al. 2017), emClarity (Himes and Zhang, 2018), and cryoSTAC (Zhang, 2019). Although most software packages support manual picking particles on orthogonal slices, it is labor-intensive and highly subjective. Automatic picking methods have been proposed to pick millions of macromolecules effectively. Automatic picking methods are divided into the reference-based method and reference-free method. The typical reference-based method is template matching (Böhm et al., 2000), which calculates cross-correlation between a predefined template and segmented volumes to find particle locations and orientations. The template can be an assembly of simple 3D shapes, a structure from Protein Data Bank, or an averaged structure from several manual samples. However, template matching still has several limitations. The quality of the predefined template has a great impact on the final matching results. The cross-correlation threshold needs human intervention. And the computation time for 3D cross-correlation increase as the types of macromolecules increase. The difference of gaussian (DoG) (Pei et al., 2016) is the most common reference-free method. This method subtracts two gaussian filtered images to find the edge of particles, but the performance of DoG highly depends on the selected gaussian filters.

In recent years, machine learning has been implemented for particle localization and classification (Chen et al., 2012). This method uses template matching results as particle candidates, calculates corresponding features for particle candidates, and classifies these particle candidates via support vector machines (SVM). Nevertheless, the limitations that existed in SVM are the dependence on template matching results and manually constructed features.

With the improvement of data acquisition for cryo-electron microscopy (cryo-EM) and Cryo-ET, deep-learning gains popularity for detecting and classifying 2D and 3D particles. Several convolutional neural networks have been designed to pick single particles in Cryo-EM images (Zhang et al., 2019; Wang et al., 2016; Zhu et al., 2017; Xiao and Yang, 2017; Bell et al., 2018). These early attempts on the 2D particle picking provide valuable insights, but they are not suitable for 3D particles and lack identifying different types of particles.

For cryo-ET data, the first deep-learning-based work is to annotate tomogram slice-by-slice using a segmentation network (Chen et al., 2017). Another deep-learning-based work is locating and identifying structures of interest per slice simultaneously using faster-rcnn (Li et al., 2019). In the above works, the tomographic reconstruction is viewed as a stack of 2D tomographic slices, and all operations are performed on slices rather than in 3D. The operations on a 2D slice are fast, and the model complexity is reduced. Nevertheless, there are limitations in slice-by-slice operations. It only focuses on the information on a single slice; hence it lacks consideration of spatial information between a set of consecutive slices. Besides, the outputs of the above networks cannot describe the particle density in a 3D tomogram precisely. Instead, 3D CNNs can pay equal attention to all directions. Especially, 3D CNNs have a better feature extraction for some small structures which does not have enough features in 2D slices. SHREC’19 challenge has indicated that deep-learning-based methods with 3D convolutions outperform those with 2D convolutions (Gubins et al., 2019) in the task of particle localization and classification.

A study uses a 3D CNN with an encoder-decoder architecture for subtomogram semantic segmentation (Liu et al., 2018). DeepFinder uses a classic 3D U-net architecture for macromolecules localization and classification in tomograms (Moebel et al., 2020). The supervised 3D CNNs methods require large amounts of training data to achieve accurate results. Nevertheless, it is hard to collect a lot of non-simulated training data in the cryo-ET research domain. SHREC challenge provides a benchmark to compare and evaluate different methods for particle localization and classification in Cryo-ET data (Gubins et al., 2019; Gubins et al., 2020). This challenge boosts the development of macromolecules detection and innovation in computational methods. In the recently launched SHREC’20 challenge, a method uses a sliding window to select a region and then classify the selected region via 3D ResNet. The sliding window process wastes lots of time because some non-particle regions are also sent to the network for inference. A method called YOPO is based on a 3D object detection network without the need for segmentation maps, but segmentation maps are important for determining the orientations of particles. Dn3DUnet method and UMC method are both based on 3D U-net with connected component analysis. The connected component analysis is a simple way to cluster the segmentation maps, while mean-shift clustering can handle more difficult situations. Methods based on adversarial learning (Lin et al., 2019), an auto-encoding classifier (Liu et al., 2019), and a 3D classification network (Che et al., 2018) provide various solutions for sub-tomogram classification after extracting the sub-tomogram. The main problem here is that the 3D classification network only receives fixed-size images, which goes against classifying particles with various sizes.

To overcome the limitations mentioned above, we designed an approach based on a 3D CNN to localize and classify macromolecules in cryo-electron tomograms. 1) To improve the localization accuracy of particles on the segmented tomogram, we combine the 3D connected component with the mean-shift clustering to yield more accurate coordinates of particles. The 3D connected component calculates rough positions of particles, and then mean-shift clustering used these positions as initial seeds for refinement. 2) To solve the imbalanced size between large and tiny particles, we use a weighted focal loss for multi-class segmentation. With the prior knowledge of the characteristics of each class, the weighted focal loss assigns different weights to the loss of each class. We can pay less attention to large particles while focusing more on small particles that are hard to classify. 3) To design the network with fewer parameters, we employ a 3D Hybrid Dilated Convolution (HDC) module in the backbone of our network. The dilated convolution with a kernel size of 3×3×3 allows for a very large receptive field while only has 3×3×3 parameters. The small size of parameters can settle the problem of training on small datasets. 4) To avoid classifying particles in a time-consuming sliding window across the tomogram, we propose a two-stage approach for fast particle detection. In the first stage, we calculate the particle coordinates from a segmented tomogram. In the second stage, the particle classification is only performed on the coordinates found by the first stage. This two-stage detection approach can save time by not classifying lots of non-particle regions.

In our experiments, we evaluated our VP-Detector on simulated tomograms. The mean-shift clustering based on 3D connected component improved the F1-score. Compared to the state-of-the-art methods, our method achieved a competitive performance on localization. We also explore the influence of layers and the size of training sets. The results demonstrate that our proposed network can train on small datasets. And our two-stage VP-Detector can offer high-speed particle detection.

The rest of the paper is organized as follows. In Section 2, we describe the overall pipeline for particle detection at the beginning (section 2.1). Then, we introduce two vital components, the 3D HDC module (section 2.2.1) and dense connectivity (section 2.2.2), to construct our network. Next, we explain the network architecture (section 2.3) and the design of the loss function (section 2.4). In Section 3, experiments on simulated tomograms are discussed in detail. Lastly, we conclude our method and future works in Section 4.

## 2 Methods

This section presents the VP-Detector algorithm for particle detection via a 3D multi-scale dense convolution network (see Figure 1). We start with an overview of our approach and outline the details of two modules: particle localization module and particle classification module (Section 2.1). And then present a 3D HDC module for multi-scale features and dense connectivity (Section 2.2). Finally, we describe the architecture of the proposed network (Section 2.3) and weighted loss function (Section 2.4).

**Figure 1.**
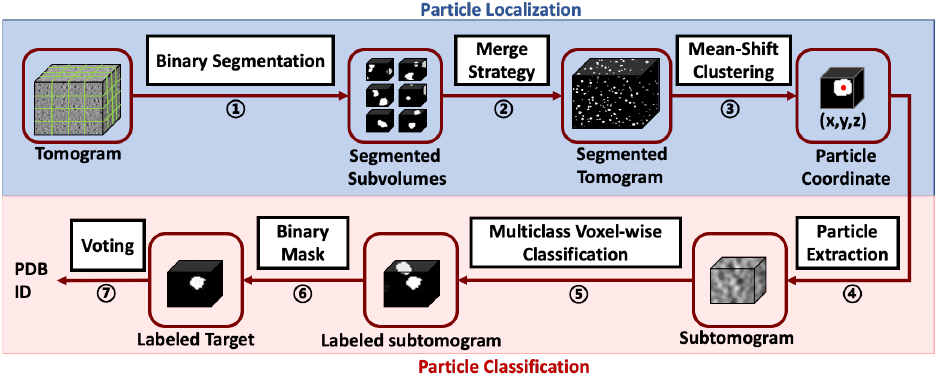
The overall workflow of VP-Detector includes seven procedures represented in white boxes. The two stages are shown in solid boxes: the particle localization stage (blue) and the particle classification stage (red).

### 2.1 Overview of the two-stage particle detector

Our particle detection algorithm provides an automatic and efficient scheme for macromolecules localization and classification in cryo-electron tomogram. Figure 1 schematically illustrates the proposed VP-Detector architecture, with seven basic procedures represented in white boxes. It is designed as a two-stage approach, including a particle localization stage (1)(2)(3) and a particle classification stage (4)–(7). Detailed descriptions of our two-stage detection are described as follows.

#### 2.1.1 Particle localization stage

The first stage conducts a fast particle localization on binary segmentation maps.

1. *Binary segmentation via a 3D MSDNet*. The goal is to obtain a binary segmentation map where each voxel is the probability of being labeled as particle class or the background class. Due to the GPU memory limit, the network conducts a binary segmentation on cubic volumes cropped from the input tomogram. In this way, we can obtain many segmented subvolumes for the following localization.
2. *Merge strategy*. All segmented subvolumes are merged into a whole segmentation map with an identical size to the original tomogram. Since cubic volumes are cropped with overlapping regions, it is rational to mask out the incomplete particles connected to the image boundary before merging them. We remove the noise in a segmentation map with the opening operation of morphology. The whole segmentation map is ready for clustering, which can determine particle localizations at one time.
3. *Clustering for finding particle coordinates*. All particles are labeled as the same value in the segmentation map, where the region for an individual particle is unclear. Ideally, each particle in a segmentation map is observed as a 3D-connected component. The connected component analysis is a frequently used method to give each component a unique label for distinguishing different particles. The rules for connected components are not rigorous enough because multiple particles connected by a few voxels can be labeled as one particle. To determine more accurate particle coordinates, we perform a clustering algorithm on the segmentation map, which can assemble voxels into groups of particles to calculate particle coordinates at sub-voxel precision. Typical clustering algorithms are k-means, mean-shift, agglomerative clustering, and DBSCAN. We use mean-shift clustering because it can handle many data points without prior knowledge of object numbers. To accelerate and improve the mean-shift clustering algorithm, we can use the centroids of 3D connected components as initial seeds for clustering. The only interactive parameter required in mean-shift clustering is the bandwidth, which is related to the particle size. Some relatively small clusters are filtered out as false positives.

#### 2.1.2 Particle classification stage

The second stage is designed to classify particles via analyzing multi-class segmentation maps.

(4) *Particle extraction*. Once we have a list of particle coordinates, we are able to extract them from the segmentation maps for the subsequent classification. Each extraction, also called a subtomogram, has a target particle at the center point.
(5) *Multi-class voxel-wise classification (segmentation) via a 3D MSDNet*. This is a vital procedure to precisely label the voxels in sub-tomogram with *N* classes. The network architecture used here is similar to that of procedure ①, where the number of layers and classes are different. Now that the labeled sub-tomograms are in place, post-processing is needed to obtain the classification results.
(6) *Masking*. The goal is to preserve the target particle only. A binary mask is utilized to wipe out irrelevant particles in the labeled subtomogram, which means only the 3D-connected component at the center is the useful information.
(7) *Voting*. The classification of a labeled target is determined by voting strategy. We count the number of class labels and taking the majority of labels as a result.

Overall, our two-stage detection approach has several advantages over one-stage detection. First, the classification only performs on the given coordinate instead of a sliding window across the segmentation map. It helps to dramatically accelerate particle detection. Second, the two-stage approach is more flexible since each stage can be used independently for different needs. For instance, the coordinates generated from the first stage can be used for testing other classification methods. Besides, the localization can deal with both known and unknown structures since the network of the first stage is trained for particle class and background class. Third, a one-stage network for localization and classification simultaneously puts a strain on hardware resources. We introduced two independent networks for localization and classification, respectively, so that each network can be more thoroughly trained under the limited hardware. Moreover, we find a precise solution to particle coordinates by combining mean-shift clustering and connected component analysis.

### 2.2 The main components of 3D MSDNet

#### 2.2.1 3D Hybrid Dilated Convolution for Multi-Scale Features

Dilated convolution is suitable for voxel-wise dense prediction because it allows enlarging the receptive field without losing resolution or coverage (Yu and Koltun, 2015). We employ 3D dilated convolutions with different dilation factors to systematically extract multi-scale features. However, the use of serialized layers with increasing dilation factor may suffer a gridding effect. To address this problem, we design an HDC module that consists of *n* subsequent convolutional layers that apply kernel size *K* × *K* × *K* with different dilation factors of [*s*_1_,…, *s_i_*,…, *s_n_*]. In our HDC module, the assignment of dilation factors follows the rule in (Wang et al., 2018). The maximum dilation factor *M_i_* in *i^th^* layer is equal to *max*[*M*_*i*+1_ − 2*s_i_*, *M*_*i*+1_ − 2(*M*_*i*+1_ − *s_i_*), *s_i_*], and we should let *M*_2_ ≤ *K* with *M_n_* = *s_i_*. Such an assignment ensures that the final size of the receptive field fully covers a square region without any holes or missing edges. Another problem that exists in dilated convolutions is that large dilation factors might capture irrelevant long-ranged information. A study has revealed that a filter with a small dilation factor can be applied to most of the valid regions on a feature map instead of the padded region (Chen et al., 2017). In our network, we use small dilation factors to capture effective multi-scale information. The number of parameters should be small, thus we adopt dilated convolutions with a kernel size of 3×3×3.

Figure 2 depicts an example of two channels HDC module with dilation factors ∈ [1,3]. HDC module applies continuous dilated convolutional layers with increasing dilation factors to effectively expand the receptive field (RF) to aggregate multi-scale contextual information. For a voxel in a feature map in *i*^th^ layer, RF_i_ represents all the voxels from feature maps in (*i* − 1)^*th*^ layer that affect its value, which will be (2*i* + 1)^3^. And *RF*′_i_ represents all the voxels from the original image that affect its value, which will be *RF*′_i_ = [(*i* + 1)*i* + 1]^3^. A number of HDC modules are grouped to form the main architecture of our network with the same pattern of dilation factors. By doing this, the dilation factors of the convolutional layers are equal to *s_i_* ≡ (*i* + 1) *mod* 4.

**Figure 2.**
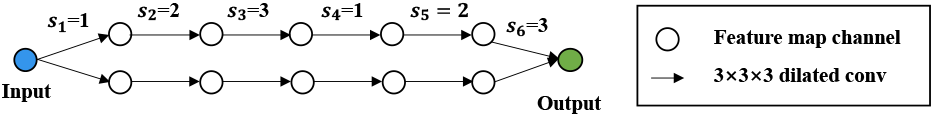
An example of two channels HDC modules with six convolutional layers with dilation factors ∈ [1,2,3].

One benefit of the HDC module is that the use of arbitrary dilation factors captures global context and detail information while preserving resolution as well as computational efficiency. Unlike other scaling operations to extract features at various scales, e.g., pooling operations or strided convolutions lose small-scale objects and detailed information. And stacking convolutional layers or increasing the filter size to acquire global contextual information could raise the number of parameters and the complexity of the model. The more important point is that our HDC module alleviates the gridding issue, which hampers the performance.

#### 2.2.2 Dense connectivity

In the densely connected network (Huang et al., 2017), each layer is directly connected to all the subsequent layers. Without loss of generality, the *i^th^* layer receives all the preceding feature maps ***Z*_0_**,…, ***Z***_*i*−1_ as inputs to generate ***Z***_i_,

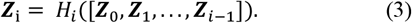

Here, [···] is an operation used to combine features, which usually uses channel-wise concatenation or element-wise addition. It introduces direct connections between any size-matched two layers. The dense connectivity has several merits: it helps alleviate the vanishing gradient problem and substantially reduces the parameters. In addition, it ensures that preceding feature maps can be fully re-used and more spatial resolution are reserved.

### 2.3 The Architecture of 3D MSDNet

In this section, we describe our proposed 3D MSDNet. Our segmentation network aims to create a label map ***y*** where voxels are labeled with categories. Suppose an input 3D image ***x*** has *C* channels, long *L*, height *H*, and width *W*. In our 3D MSDNet, the *i^th^* layer outputs a feature map 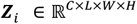, and a single channel *j* of ***Z**_i_* is denoted as 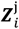.

#### Composite function for convolutional layers

We define 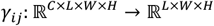 as a function to convolve the input feature map in *i^th^* layer with different kernels for each channel and sums up the resulting images voxel by voxel to yield a single channel *j* of the output feature map,

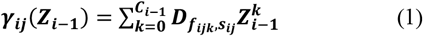

where D_***f***_***Z*** is a 3D convolution of image ***Z*** with filter ***f***. During convolution operations, dilation factors s for each channel of a certain layer are different. And ***f**_ijk_* are kernels of size 3 × 3 × 3 voxels that have fewer parameters and a more enhanced capability of nonlinear mapping than larger kernels. *C*_*i*−1_ is the number of channels of an output feature map in *i* − 1^*th*^ layer.

Convolutional layers have a composite operation *H*: performing a 3D convolution on each channel of the input with a different filter, summing up the resulted images voxel by voxel, adding a bias, and applying an element-wise nonlinearity. Thus, the single channel output 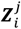 of such a convolutional layer is defined as,

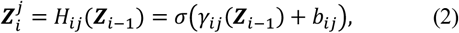

where 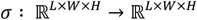 denotes an activation function, such as sigmoid, rectified linear units (ReLU), softplus, leaky ReLU (LReLU). And 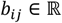 is the bias for channel *j* of an output feature map in *i^th^* layer.

#### First layer

The input 3D image ***x*** is taken as the first layer ***z*_0_** and is defined as a set of voxels 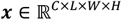. The single channel *j* of ***x*** is denoted as ***x**^j^*.

#### Subsequent layers

In the HDC mentioned above module with padded convolutions, the input image, all feature maps, and output image have identical dimensions. Therefore, all previously computed feature maps can be used to calculate the feature map in current layer. To further enhance the information flow between layers, we introduce dense connectivity to directly connect any layer to all subsequent layers (Huang et al., 2017). Here, the *i^th^* layer receives all the preceding feature maps ***Z***_0_,…, ***Z**_*i*−1_* as input, thus a single channel output feature map 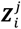 is defined as:

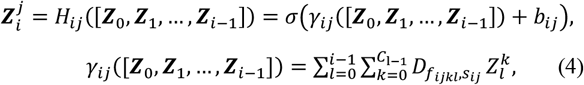

where *H_ij_* is a composite operation for channel *j* of the output in *i^th^* layer, and *σ* is an activation function. We adopt LReLU as the activation function, that is *f*(*x*) = *max*(0.01*x,x*), to avoid the dying ReLU problem. LReLU brings the tradeoff between the network sparsity and the performance and solves the vanishing gradient problem (Zhang et al., 2017). In equation 4, *γ_ij_*([***Z***_0_, ***Z***_1_,…, ***Z***_*i*−1_) is an operation of convolving all previously computed feature maps from the first layer to *i* − 1^*th*^ layer and summing up the resulting images voxel by voxel to yield a single channel *j* of output feature map in *i^th^* layer. And ***f**_ijk_* are kernels of size 3 × 3 × 3 voxels. The dense connectivity utilizes element-wise addition, defined as [···], to aggregate each feature map within the network. Element-wise addition has several merits: Its summation operation accelerates the training process via parallelization as well as dense connectivity. This indirectly helps solve the vanishing gradient problem. Besides, it supports incorporating feature maps with various size receptive fields into a feature map with new features. Compared to concatenation, element-wise addition performs better on small and blurry objects in images (Gao et al., 2018). In our research, since the particles have low signal-to-noise (SNR) ratio caused by low electron doses, the contextual information is useful for their detection. Element-wise addition is strongly recommended because it can learn the relationship between the target and context well while concatenation cannot. Our 3D MSDNet allows that the feature map per layer has the different number of channels. For simplicity, suppose the number of channels for all layers is a fixed value, then we can define the number of non-input and non-output layers as the depth *d* and the number of channels as the width *w*. Figure 3 illustrates the layout of the 3D MSDNet with *w* = 2 and *d* = 3.

**Figure 3.**
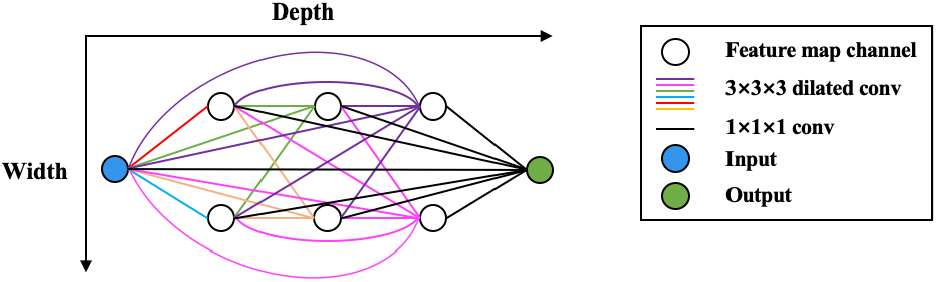
An illustration of MS-D network with *w* = 2 and *d* = 3. Colored lines denote dilated convolutions with different dilation factors.

#### Last layer

The final layer output an image ***y*** through similar composite operations and the channel *j* of ***y*** is defined as,

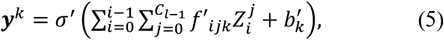

where *f′_ijk_* are kernels of size 1 × 1 × 1 voxels. It computes a linear combination of feature maps of the input layer and intermediate layers. The nonlinear mapping *σ′* is a voxel-wise soft-max function for dense image labeling. Our network gives a voxel-to-voxel prediction so that the sizes of the output image and input image are identical.

#### Parameters

The trainable parameters of our proposed network include convolution filters ***f**_ijk_*, biases *b_ij_* of non-input layers and non-output layers (see equation 4), as well as weights *f′_ijk_*, biases 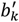 of the last layer (see equation 5). The number of filters, weights and bias are denoted as *N_f_, N_w_, N_b_* respectively, hence the number of parameters is *N_Θ_* = *N_f_ + N_w_ + N_b_*. Suppose an input image with *C* channels, we can have 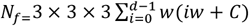, *N_w=_* (*wd* + *C*)*C*, *N_b_* = *wd* + *C*, where *d* and *w* are the depth and width of the network.

### 2.4 Weighted focal loss of 3D MSDNet

Class imbalance is a common problem in segmentation tasks where the number of voxels labeled for each class is disproportionate. In our simulated dataset, the volumetric ratio between the largest and the smallest particles can reach nearly 30:1. The loss is dominated by prevalent labels of large particle class, which leads to inaccurate labeling on small particle class. Focal loss is a modified version of cross-entropy loss by downweighting the loss assigned to easy examples (Lin et al., 2017). To address the class imbalance problem, we designed a weighted focal loss for multiclass segmentation. We defined the weighted focal loss as:

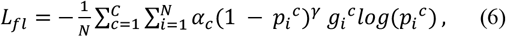

where *N* is the number of voxels in an image, *c* ∈ *C* represents a class *c, g_i_^c^* is a binary indicator of ground-truth class *c* of voxel *i*, and *p_i_^c^* represents the corresponding model’s estimated probability. *α_c_* is introduced to reweight different classes, which are based on the effective radius of particles. *γ* is a focusing parameter that down-weights the loss assigned to well-classified samples and makes hard samples contribute more to the loss. When *γ* = 0 and *α_c_* = 1, the function behaves as cross-entropy.

## 3 Results

### 3.1 Dataset

To evaluate the performance of localization and classification for biological particles in the cryo-electron tomogram, we carried out the experiments on realistically simulated tomograms provided by SHREC. Figure 4(A) gives an example of a raw tomogram. It is tough for a human to see particles inside the noisy tomogram. SHREC’20 contains nine sets of 512 × 512 × 512 tomograms with 1 nm/voxel resolution. Ground truth volumes and particle location are also given in the dataset. Between 2400 and 2800 bio-particles of 12 classes are placed in each tomogram. In our experiments, tomograms from 1 to 7 make up training sets, and tomogram 8 is for validation. Tomogram 9 is the test tomogram to evaluate our particle detector. SHREC’19 contains ten sets of tomograms generated in a similar way which has higher SNR.

**Figure 4.**
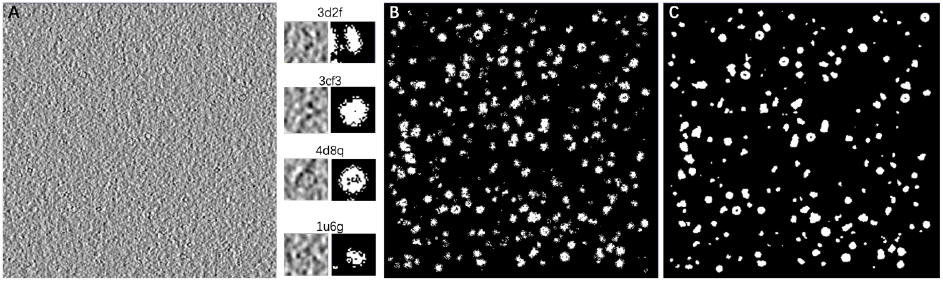
An example of SHREC’20 dataset. (A) A slice of a raw tomogram. (B) The corresponding ground truth. (C) Predicted segmentation map from our VP-Detector. 3d2f, 3cf3, 4d8q, 1u6g are four particles extracted from the raw tomogram and ground truth.

### 3.2 Implementation details

VP-Detector is built on a deep learning framework called PyTorch with CUDA acceleration, written in Python programming language. All experiments are carried on a workstation with four GeForce RTX 2080 Ti. We train our weights using stochastic gradient descent with the Adam optimizer. The initial learning rate is 0.001 and decreased by ten times after 80 epochs. We stop training after 500 epochs. The localization and classification networks are based on 3D MSDNet with different hyperparameters. The localization network has dilation factors ∈ [1,7] and 49 dilated layers with 132K trainable parameters. The classification network has dilation factors ∈ [1,5] and 100 dilated layers with 551K trainable parameters.

### 3.3 The localization performance of VP-Detector

#### 3.3.1 Evaluation Metrics

The evaluation metrics for particle localization are precision *P*, recall *R*, miss rate *M*, F1-score, and confusion matrix.

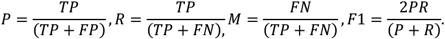

The evaluation metrics for semantic segmentation are: voxel accuracy VA, mean voxel accuracy MVA, and mean IoU MIoU. We assume that *P_ij_* is the number of voxels of class *i* predicted to belong to class *j*, and the number of *M* classes.

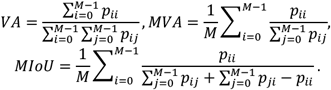

#### 3.3.2 Qualitative Evaluation

We compare VP-Detector with all algorithms submitted to SHREC’19 and SHREC’20 challenge, and the performance comparison is listed in Table 1,2. The best results are shown in bold. For the SHREC’20 dataset, table 1 indicates that VP-Detector has the best precision of 0.978 and the best F1-score of 0.951. For the SHREC’19 dataset, Table 2 shows that VP-Detector has a superior performance and has the best evaluation metrics of recall, miss rate, F1-score of 0.86299, 0.13701, and 0.88709, respectively. Precision is the ratio of correctly detected particles to the detected results. Precision is an essential metric for particle localization because the false positives interfere with the subsequent classification, alignment, and averaging structures. F1-score is a weighted average of precision and recall. VP-Detector is proved to be the most well-perform among all localization methods since it has the best F1-score.

**Table 1.**
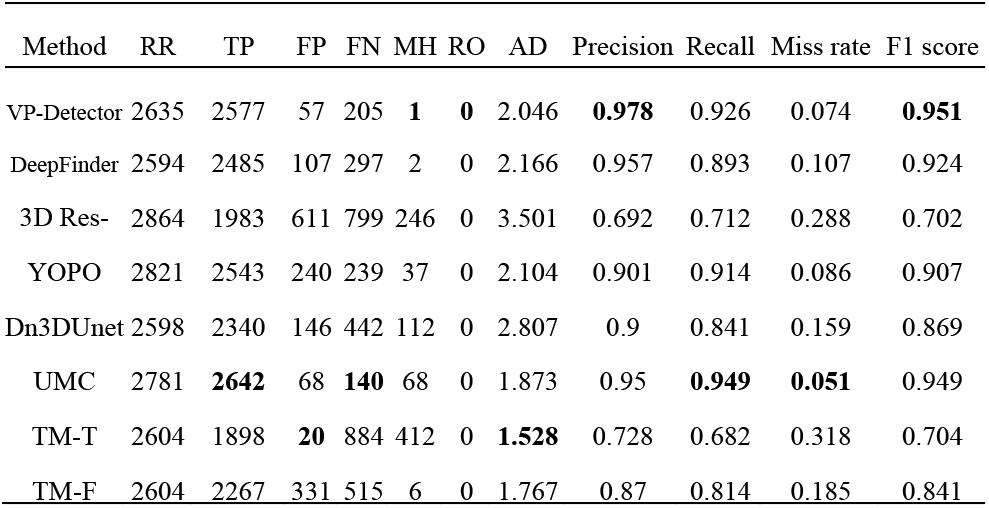
Performance comparison between different algorithms for the SHREC’20 dataset. RR (reported results), MH (unique particles that had duplicate results), RO (results not inside particle region), AD (average Euclidean distance between the predicted center and ground truth center).

| Method | RR | TP | FP | FN | MH | RO | AD | Precision | Recall | Miss rate | F1 score |
| --- | --- | --- | --- | --- | --- | --- | --- | --- | --- | --- | --- |
| VP-Detector | 2635 | 2577 | 57 | 205 | **1** | **0** | 2.046 | **0.978** | 0.926 | 0.074 | **0.951** |
| DeepFinder | 2594 | 2485 | 107 | 297 | 2 | 0 | 2.166 | 0.957 | 0.893 | 0.107 | 0.924 |
| 3D Res- | 2864 | 1983 | 611 | 799 | 246 | 0 | 3.501 | 0.692 | 0.712 | 0.288 | 0.702 |
| YOPO | 2821 | 2543 | 240 | 239 | 37 | 0 | 2.104 | 0.901 | 0.914 | 0.086 | 0.907 |
| Dn3DUnet | 2598 | 2340 | 146 | 442 | 112 | 0 | 2.807 | 0.9 | 0.841 | 0.159 | 0.869 |
| UMC | 2781 | **2642** | 68 | **140** | 68 | 0 | 1.873 | 0.95 | **0.949** | **0.051** | 0.949 |
| TM-T | 2604 | 1898 | **20** | 884 | 412 | 0 | **1.528** | 0.728 | 0.682 | 0.318 | 0.704 |
| TM-F | 2604 | 2267 | 331 | 515 | 6 | 0 | 1.767 | 0.87 | 0.814 | 0.185 | 0.841 |

**Table 2.**
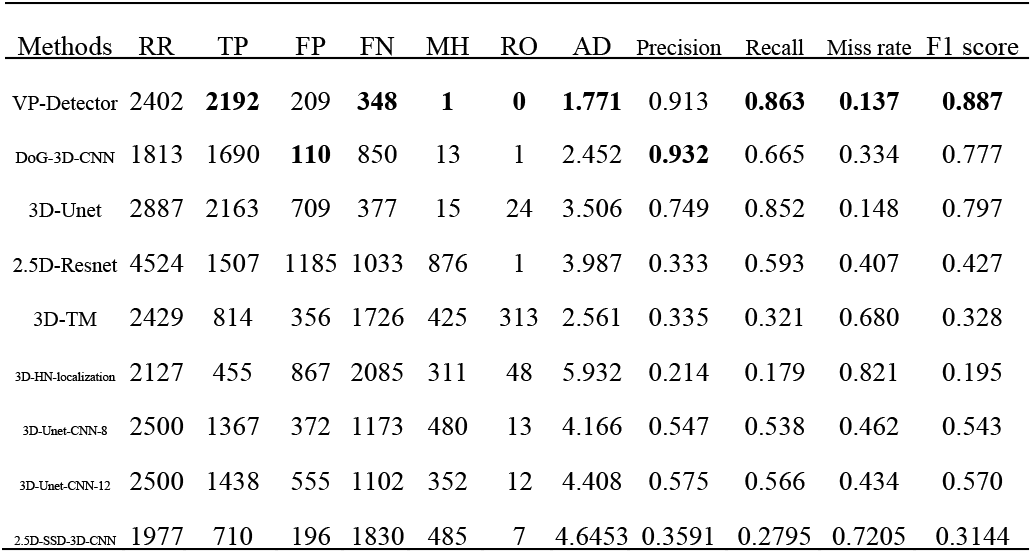
Performance comparison between different algorithms for the SHREC’19 dataset.

We use mean-shift clustering to improve the performance of localization. Before mean-shift clustering, 87 particles had more than one prediction. The precision and F1-score are 0.916 and 0.931. After mean-shift clustering, there is one particle that had more than one prediction. The precision improves 0.062, and F1-score improves 0.02.

#### 3.3.3 Visualization

Figure 4(C) shows the segmentation map of the input tomogram generated by our VP-Detector. Our VP-Detector shows excellent robustness to the noise. We can also observe that particles on the prediction map have smoother and clearer boundaries than ground truth volume shown in Figure 4(B). We use VA, MVA, and MIoU to prove the effectiveness of our prediction map with ground truth volume. Here, VA is 0.961, MVA is 0.819, and MIoU is 0.763.

### 3.4 The classification performance of VP-Detector

#### 3.4.1 Influence of loss function

We trained a classification network with cross-entropy loss and a classification network with weighted focal loss. As plotted in Figure 5, all losses decline sharply in the first several epochs and become steady during more epochs. The weighted focal loss is much lower than cross-entropy loss. We can also observe that the validation loss of weighted focal loss converges earlier than cross-entropy. However, the validation loss of crossentropy presents a slight shaking. Moreover, we evaluate the class accuracy for cross-entropy loss and weighted focal loss. The results are presented in Figure 6, where the accuracy for a certain PDB is defined as correctly found particles / the total number of particles. The top figure depicts the class accuracy for cross-entropy loss. Most of the classes are not stable during training, and they tend to be up and down many times. It can only classify large particle class well. And it is hard to pick the best-fit model for all classes. The bottom figure depicts the class accuracy for weighted focal loss. All the classes become steady after 250 epochs. Table 3 compares 100 layers of 3D MSDNet with cross-entropy loss and weighted focal loss. Particles are grouped into four sets according to sizes. The network with weighted focal loss has the highest class accuracy for small, medium, large particles.

**Figure 5.**
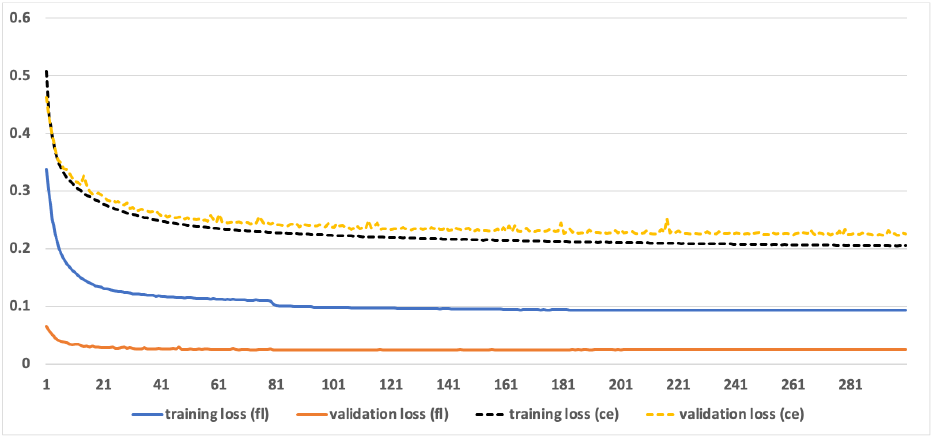
The Training loss and validation loss for different loss functions.

**Figure 6.**
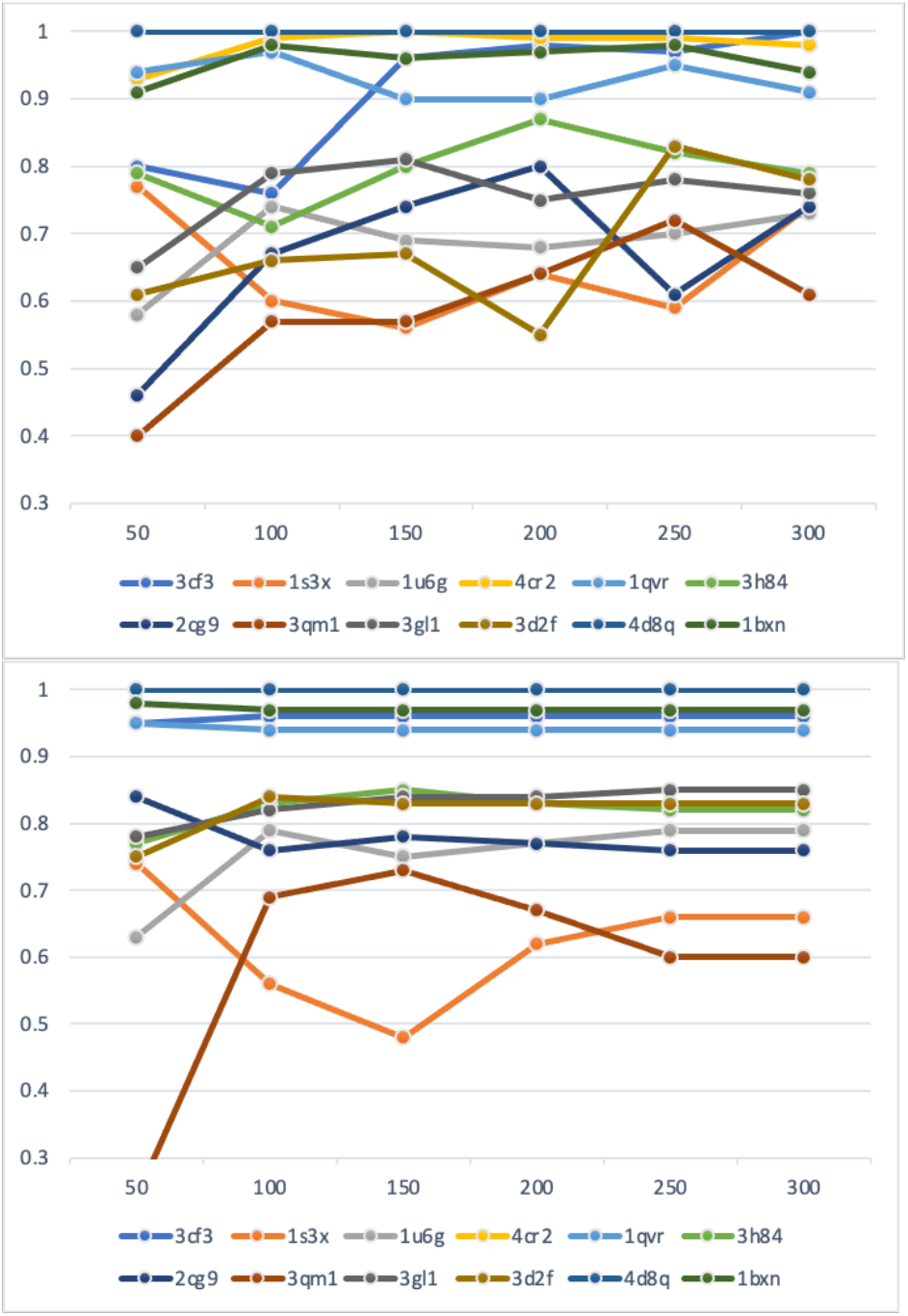
The accuracy of 12 classes for different loss functions. Top figure uses cross-entropy loss function. Bottom figure uses weighted focal loss.

**Table 3.**
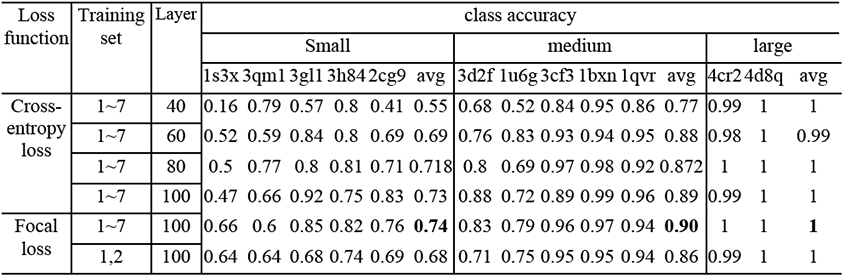
Performance of classification with different hyperparameters for SHREC’20 dataset.

The weighted focal loss function performs better than the cross-entropy loss function in terms of loss and class accuracy during training. The crossentropy loss is dominated by prevalent labels of large particle class, which lacks training on small particle class. In contrast, weighted focal loss can pay more attention to hard classes by adding a weight factor. Each class can be fully learned during training, no matter how small. The classification confusion matric of our VP-Detector is offered as shown in Figure 7. Although we add more weight to the loss of small classes, it is difficult for our network to identify between 1s3x and 3qm1. Sometimes our network may confuse 2cg9 with 3gl1, 3h84 with 3d2f, and 1u6g with 2cg9. We find that the confused pair have a similar effective radius. The high noise of tomograms increases difficulty for confused pairs.

**Figure 7.**
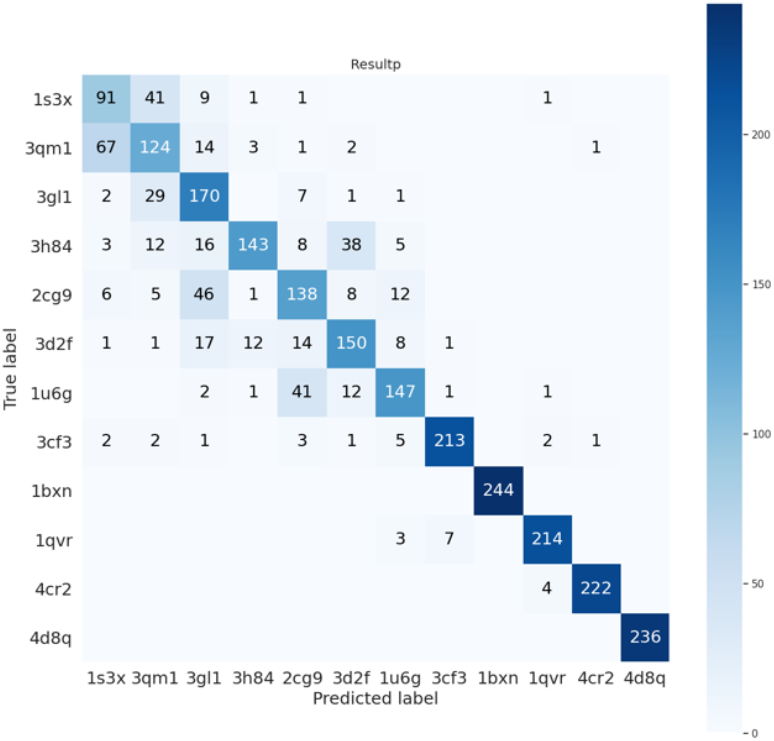
Classification confusion matric of our VP-Detector.

#### 3.4.2 Influence of hyperparameters

The classification result on a test tomogram is presented in Table 3. We discuss the influence of layers and the sizes of training sets. The number of layers is not sensitive to three classes (1bxn, 4d8q, 4cr2), and shallow networks can recognize them effectively. 1s3x and 3qm1are the hardest classes for both shallow and deep networks. We divide 12 classes into four groups based on their difficulty, as shown in Table 4. We find that the classification difficulty depends on the effective radius, *R_e_* = 3*V/A*, where *V* is the volume and *A* is the surface area. The smaller the effective radius, the harder it is to identify.

**Table 4.** The effective radius of 12 classes. The classes colored in orange are very difficult. The classes colored in yellow are relatively difficult. The classes colored in green are normal. The classes without color are easy.

| PDB | 1s3x | 3qm1 | 3h84 | 1u6g | 3d2f | 2cg9 | 3gl1 | 1qvr | 3cf3 | 4cr2 | 1bxn | 4d8q |
| --- | --- | --- | --- | --- | --- | --- | --- | --- | --- | --- | --- | --- |
| $R_e$ | 2.56 | 2.88 | 2.822 | 3.065 | 3.145 | 3.108 | 3.065 | 3.248 | 4.181 | 4.696 | 4.78 | 4.85 |

There are seven sets of tomograms for us to prepare the training sets. When we use only two sets of tomograms for training, the large classes have almost the same accuracy, and the small classes and medium classes have a slight decline. In this way, our proposed 3D MSDNet supports training on small datasets since we combine HDC module and dense connection to design the network with fewer parameters.

### 3.5 Run Time for inference

Table 5 shows the run time for different methods. The test tomogram has a size of 512 × 512 × 512 voxels with 2782 particles. The traditional template matching methods are the most time-consuming methods, which cost a few hours. The CNNs based method using slide window needs about an hour or more to finish particle detection. Our method uses only 13 minutes 44 seconds to achieve accurate particle detection, which creates a balance between efficiency and accuracy. Our fast detection has two stages. The localization stage costs about 8 minutes, and the classification stage costs about 5 minutes. Our two-stage detection approach is fast because the classification is only performed on given positions.

**Table 5.** Run time for particle detection on a test tomogram.

| Method | Inference time |
| --- | --- |
| 3D MSDNet | 13m44s = 8m16s(localization) + 5m28s(classification) |
| DeepFinder | 20m |
| 3D ResNet | 2h |
| YOPO | 40m |
| Dn3DUnet | 1m 41s |
| UMC | 42m |
| TM-T | 27h 24m |
| TM-F | 1h 24m |

## 4 Discussion

In this paper, we present an automatic and accurate CNNs based approach for volumetric particle detection in cryo-electron tomogram. To solve the problem of training on relatively small datasets, we design a novel 3D convolutional network architecture with HDC modules and dense connectivity. The 3D HDC module is a set of dilated convolutional layers with various dilation factors to capture multi-scale objects and contextual information. Dense connectivity enhances the re-use of feature maps. We evaluated the VP-Detector on the simulated tomograms. Comparing to the state-of-the-art methods, our method achieved the best localization performance with the highest F1-score. The mean shift clustering also helps a better F1-score. Experimental results also demonstrate that our network can train on small datasets. Moreover, our two-stage detection approach can achieve fast detection for about 14 minutes. In future work, we plan to improve memory efficiency by sharing the memory of the same features (Pleiss et al., 2017). Besides, there is still room for improvement towards small object detection. We can explore more future fusion techniques focusing on detailed information. In addition, preprocessing for input tomograms may improve the performance of particle detection, such as image denoising (Zhang et al., 2017).

